# *SELF PRUNING 3C* is a flowering repressor that modulates seed germination, root architecture and drought responses

**DOI:** 10.1101/2022.03.10.483842

**Authors:** Juliene dos Reis Moreira, Alejandra Quiñones, Bruno Silvestre Lira, Jessenia M. Robledo, Shaun J. Curtin, Mateus H. Vicente, Dimas M. Ribeiro, Malgorzata Ryngajllo, José M. Jiménez-Gómez, Lázaro Eustáquio Pereira Peres, Maria Magdalena Rossi, Agustín Zsögön

## Abstract

Allelic variation in the *CETS* (*CENTRORADIALIS, TERMINAL FLOWER 1, SELF PRUNING*) gene family has been shown to control agronomically important traits in many crops. *CETS* genes encode phosphatidylethanolamine binding proteins (PEBPs) that have a central role in flowering time control as florigenic and anti-florigenic signals. The great expansion of *CETS* genes in many species suggests that the functions of this family go beyond flowering. Here, we characterize the tomato *SELF PRUNING 3C* (*SP3C*) gene, and show that besides acting as a flowering repressor it also regulates seed germination and modulates root architecture. We show that loss of *SP3C* function in CRISPR/Cas9-generated mutant lines accelerates seed germination and increases root length with lower root side branching. Higher *SP3C* expression in transgenic lines promotes the opposite effects and also improves tolerance to water stress in seedlings. These discoveries provide insights into the role of *SP* paralogs in agronomically relevant traits and support future exploration of the involvement of *CETS* genes in abiotic stress responses.

**Highlight:** The *SELF PRUNING 3C* (*SP3C*) gene is a repressor of flowering in tomato and exhibits additional functions, acting as a repressor of seed germination and modulating root architecture.

## Introduction

Domestication followed by breeding practices allowed the selection and improvement of traits of agricultural interest in most crops. A common set of traits were the target of human selection, including fruit size, colour and shape, along with changes in shoot architecture, alterations in flowering time, reduction of seed dispersal and dormancy and modifications in roots (Bai and Lindhout, 2007; Meyer and Purugganan, 2013). The ‘domestication syndrome’ comprises changes in these traits and underpins the adaptation of crops to agricultural settings, generally in monoculture, which demands large-scale production and cultivation under different environmental conditions (Denham *et al*., 2020). A deeper understanding of the genetic control of domestication traits could allow the targeted manipulation of genomes using gene editing tools to rapidly create tailored agronomic traits (Zsögön *et al*., 2017).

Some of the genes that control domestication and breeding-targeted identified in wheat, sugar beet, sunflower, barley, rice, potato and tomato include allelic variation in the coding or regulatory sequences in members of the *CETS (CENTRORADIALIS* in *Antirrhinum majus; TERMINAL FLOWER 1* in *Arabidopsis thaliana; SELF PRUNING* in *Solanum lycopersicum*) gene family (Bonnin *et al*., 2008; Pin *et al*., 2010; Blackman *et al*., 2010; Comadran *et al*., 2012; Ogiso-Tanaka *et al*., 2013; Kloosterman *et al*., 2013; Soyk *et al*., 2017). *CETS* genes encode Phosphatidylethanolamine Binding Proteins (PEBPs) and play a central role in flowering control (Wickland and Hanzawa, 2015; Susila *et al*., 2021). They are grouped into three clades, classified according to the sequence similarity with the founding members in *Arabidopsis: FLOWERING LOCUS T* (*FT*-like flowering inducers, *TERMINAL FLOWER1* (*TFL1*)-like flowering repressors, and *MOTHER OF FT AND TFL1* (*MFT*-like, an ancestral clade without clear effects on flowering, but with other demonstrated effects on seed germination (Chardon and Damerval, 2005; Hedman *et al*., 2009; Yu *et al*., 2019).

The expansion of the *CETS* gene family in tomato, *Solanum lycopersicum*, resulted in 12 paralogues distributed into seven chromosomes, with five grouped as *FT*-like, five as *TFL1*-like and two as *MFT*-like (Lifschitz *et al*., 2014; Cao *et al*., 2016). A mutation in the coding region of the *SP* gene determines a conversion from indeterminate to determinate shoot growth via changes in auxin signaling (Silva *et al*., 2018), with a greater number of inflorescences and highly synchronized fruit ripening. The *sp* mutation revolutionized tomato cultivation and is extensively used in breeding programs for the selection of industrial tomato cultivars, resulting in compact and productive plants that allow high density field cultivation and mechanized harvesting (Park *et al*., 2014). Another *CETS* paralogue, *SELF-PRUNING 5G* (*SP5G*) acts as a flowering repressor under long-day conditions, and a spontaneous mutation in the *cis-* regulatory region was selected to produce photoperiod neutral tomato plants (Soyk *et al*., 2017; Zhang *et al*., 2018). Changes in the balance of anti-florigenic *SP* and florigenic *SINGLE FLOWER TRUSS* - *SFT* alleles generate heterosis for yield (Krieger *et al*., 2010) by altering the vegetative to reproductive growth balance (Vicente *et al*., 2015). The manipulation of the *SP*/*SFT* balance using targeted gene editing tools allows the tailoring of shoot architecture to optimize plant growth and yield (Jiang *et al*., 2013; Rodríguez-Leal *et al*., 2017).

The manipulation of *CETS* genes in tomato and other crops has allowed improvement of elite varieties (McGarry and Ayre, 2021) or even domestication of wild species through directed genetic editing (Zsögön *et al*., 2018). However, other traits besides those of the canonical ‘domestication syndrome’ may have been targeted, and may include altered responses to stress conditions (Milla *et al*., 2015). Some roles for *CETS* members in responses to abiotic stress have been demonstrated. In *Arabidopsis, MFT* regulates seed germination under high salinity by modulating ABA signalling, and *BROTHER OF FT* (*BFT*) promotes delay in flowering under high salinity, providing an additional pathway to control flowering under salt stress (Xi *et al*., 2010; Ryu *et al*., 2011, 2014). The tomato florigen (*SFT*) is involved in control of stomatal opening (Kinoshita *et al*., 2011) and regulates water-use efficiency (Robledo *et al*., 2020).

Considering the large expansion of *CETS* in many species and evidence that these genes can regulate important physiological processes under stressful environmental conditions, the study of *CETS* genes is an important tool for improvement of crops in the face of challenging conditions for agriculture in the coming years (Zsögön *et al*., 2021). Here, we characterized *SELF PRUNING 3C* (*SP3C*), a member of the tomato *CETS* family whose function in flowering and plant development remained hitherto unexplored. We used reverse genetic approaches to create overexpressing and loss-of-function alleles for *SP3C* and evaluated its role on plant development. We demonstrated that *SP3C* acts as a flowering repressor. Transgenic plants overexpressing *SP3C* exhibited delayed flowering and increased number of leaves to anthesis. Additionally, we found that loss of *SP3C* function accelerated seed germination and increased root length but with lower root side branching, while *SP3C* overexpressing promoted the opposite effects. We also showed that *SP3C* expression is upregulated during a period of progressive drought, and the high levels of *SP3C* in overexpression lines conferred greater resistance to drought in the seedling stage.

## Materials and methods

### Growth conditions and plant material

Tomato (*Solanum lycopersicum* L.) plants, cultivar Micro-Tom (MT) (Martí *et al*., 2006) were grown in a greenhouse at Viçosa (642 m above sea level, 20°45′S, 42°51′W), Minas Gerais, Brazil, under semi-controlled conditions: mean temperature of 26/18°C day/night, photoperiod 12-h/13-h (winter/summer), ~800 μmol m^-2^ s^-1^ PAR, and irrigation to field capacity once per day. Seeds were germinated in polyethylene trays containing Tropostrato^®^ commercial substrate until development of the first true leaf. At this stage, seedlings of each genotype were transplanted to pots with capacity of 350 ml, containing substrate described above. The basic fertilizing was done with NPK 2 g L^-1^ (10-10-10) and limestone 4 g L^-1^. Two overexpressing MT lines (*35S::SP3C*#1 and *35S::SP3C*#3) and three knockout lines (*sp3c*#12, *sp3c#*21 and *sp3c*#24) were selected at the T_3_ or subsequent generations to perform the analyses.

### Phylogenetic inference and sequence analyses

The *CETS* gene sequences of domesticated tomato (cv. Heinz 1706) and its putative ancestral species *S. pimpinellifolium* (LA1589) we retrieved from the Sol Genomics Network (SGN, solgenomics.net) database. We aligned the sequences with MUSCLE and then submitted the alignment to trimA1 for alignment trimming and to FastTree for tree inference. The tree for the tomato genes was constructed using FigTree (tree.bio.ed.ac.uk/software/figtree/) and the clades were annotated using previously described phylogenetic information from the model species *Arabidopsis thaliana* (Kobayashi *et al*., 1999; Mimida *et al*., 2001; Hedman *et al*., 2009). The positions of exons and introns were retrieved from SGN and the sequence differences between species were annotated manually, the gene alignments were performed and the figure drawn with Geneious R9.

Protein sequences were retrieved from Phytozome v12.1 database by searching the families containing putative CETS (CENTRORADIALIS/TERMINAL FLOWER 1/SELF-PRUNING)-like proteins described in Cao *et al*., 2016. Most proteins were clustered in family 92441556 and a few in family 92426874 and 92423510. All sequences from *Arabidopsis thaliana*, *Solanum lycopersicum*, *Coccomyxa subellipsoidea* C-169 and *Dunaliella salina* were retrieved (Supplementary table S1) and aligned using the M-Coffee algorithm (Wallace *et al*., 2006). The phylogenetic tree was reconstructed from the alignment using the PhyML 3;0 package (Guindon *et al*., 2010), using the LG substitution model, tree optimization by tree topology and branch length, tree improvement by subtree pruning and regrafting, and branch support by SH-like analysis (www.hiv.lanl.gov/content/sequence/PHYML/interface.html). Chlorophyta species were used as outgroup. *S. lycopersicum* proteins were named accordingly to the chromosome and bin position described in (Eshed and Zamir, 1995).

For promoter analysis, the sequences of *SP*, *SFT* and *SP3C* genes (1000 bp upstream of the start codon) from *S. lycopersicum* cv. Heinz (1706) was extracted from Sol Genomics database (solgenomics.net/, ITAG release 2.40). Putative transcription factors binding sites and their role in biological processes were identified using the PlantCARE database (Lescot *et al*., 2002).

### Gene expression analyses

Tomato plants, *S. lycopersicum* cv. M82 and *S. pennellii* LA716 were grown from seeds in square pots (7 × 7 × 8 cm) filled with a standard soil (Einheitserde Special, Typ Mini Tray, Einheitserde- und Humuswerke Bebr. Patzer, Sinntal Altengronau, Germany) mixed with fertilizer (Osmocte Start, 111-11-17 2MgO + TE, Everris Start) in ratio of 70 L of soil to 100 mL of the fertilizer. Plants were kept in a growth cabinet (Conviron) at 23/20C, with 55/65% relative humidity and 16h/8h light/dark periods, and light intensity of 130 μmol m^-2^ s^-1^. At the time of sampling *S. lycopersicum* seedlings had 2 compound leaves and *S. pennellii* had 3 small compound leaves and a leaf bud.

Throughout growth, the water content (WC) of the soil was maintained at 56% by daily rewatering. Added water was calculated by weighing of the plants. Drought was introduced after two weeks of growth by halting watering in the drought-treated plants. Plants were collected at the 5^th^, 8^th^ and 11^th^ day of drought treatment, which corresponds to decreasing soil water content in 40%, 30%, 20%, respectively. At each time point, compound leaves of 3 plants per condition and genotype were harvested and flash frozen in liquid nitrogen. Total RNA was independently extracted from 40 mg of grounded leaves of each plant using RNeasy Plant Mini Kit (Qiagen, Hilden, Germany). Samples were treated on column with DNase using RNase-Free DNase Set (Qiagen, Hilden Germany). Libraries were prepared according to the Illumina TruSeq RNA protocol and sequenced on an Illumina HiSeq2500 machine (Illumina, Inc., San Diego, CA) at the Genome Center of the Max Planck Institute for Plant Breeding Research Cologne.

We obtained a total of 423.91 million 100bp single end reads (average 11,8 million reads per sample, median 12.4 million reads per sample) that were aligned to the tomato genome reference sequence v4.0 using HISAT2 v 2.1.0 (Kim *et al*., 2019) with default parameters except for maximum intron length, that was set to 218033. An average of 90,5% of the reads were aligned to the reference (96,5% for M82 reads and 83,9% for *S. pennellii* reads). The number of reads per transcript was counted with the feature Counts function in the Rsubread package in R (Liao *et al*., 2019). We surveyed the homogeneity of the samples with the PoissonDistance function in the Bioconductor PoiClaClu package (Witten, 2011). RPKMs were calculated for each gene and sample by dividing the number of reads by the length of the gene in kilobases and by the number of millions of reads in each sample. Short reads in this project have been deposited at SRA under project number PRJNA800740.

### Molecular cloning and plant transformation

The coding sequence of the *SP3C* gene (528 pb) was amplified by Polymerase Chain Reaction (PCR) from cDNA extracted from leaves of *Solanum lycopersicum* cv. Micro-Tom, purified and cloned in the plasmids pCR8/GW/TOPO^®^ (Invitrogen) using the primers listed in Supplementary Table S2. Subsequently, the fragments were combined into the overexpression vector pK7WG2D by Gateway^®^ technology (Karimi *et al*., 2007). The correct assembly of the construct was confirmed by PCR and Sanger sequencing and used for *Agrobacterium tumefaciens* (EHA105 strain) transformation by thermal shock.

To generate a knockout in the *SP3C* gene, a CRISPR/Cas9 vector containing two guide RNAs (gRNAs) were designed to direct the Cas9 enzyme to two coding regions in the first and fourth exon. The gRNAs 5′-CAACCTCGCGTCGAAATTGG-3′ and 5′-TGGATCCCAATTACCGGCTT-3′ were artificially synthesized and cloned into the pMDC32 vector (Curtis and Grossniklaus, 2003) by a Golden Gate reaction (Weber *et al*., 2011). The plasmids were confirmed by Sanger sequencing and then used for *Agrobacterium tumefaciens* (GV3101 strain) transformation by thermal shock. Both target sites contained restriction enzyme sites, which facilitated mutant screening by PCR followed by endonuclease digest assays. We ensured that the designed gRNAs would not target other genes by searching for potential off-targets using the Cas-OFFinder algorithm against the tomato genome assembly (solgenomics.net/, ITAG release 4.0) (Bae *et al*., 2014).

For tomato *Agrobacterium tumefaciens-mediated* transformation, the EHA105 (overexpression vector) and GV3101 (CRISPR/Cas9 vector) strains were used according to Pino *et al*., (2010), with some modifications. Briefly, seeds were sterilized and inoculated into flasks containing MS salts (Murashige and Skoog, 1962) half-strength, vitamins B5, 30 g L^-1^ sucrose and 4 g L^-1^ agar. The pH of the medium was adjusted to 5.7 ± 0.05 and autoclaved for sterilization. Micro-Tom (MT) cotyledons from 8-days-old seedlings were sectioned and used for co-cultivation with *A. tumefaciens* in Root Inducing Medium (RIM) for two days, 25 ± 1 °C in the dark. The explants were then transferred to Shoot Inducing Medium (SIM) and kept in 25 ± 1 °C under 16/8 h photoperiod and 10-20 μmol m^-2^ s^-1^ irradiance for four weeks. Well-developed shoots (≥ 5 cm) were transferred into flasks containing MS medium without hormones, for shoot development and rooting. In all *in vitro* stages, the medium was supplemented with 300 mg L^-1^ timentin and 100 mg L^-1^ kanamycin or 6 mg L^-1^ hygromycin for selection of overexpression and CRISPR/Cas9 events, respectively.

### Selection of overexpressing and mutant lines

Regenerated overexpressing and knockout T_0_ plants were acclimatized under controlled conditions, 16/8 h photoperiod, 150 μmol m^-2^ s^-1^ irradiance and 25 ± 1 °C temperature, using autoclaved Tropostrato^®^ substrate and vermiculite (1:1) for four weeks. Further, T_0_ transgenic plants were transferred to greenhouse conditions for genotyping and seed production.

To verify the transgene insertion, genomic DNA was extracted from young leaflets as described by Dellaporta *et al*. (1983). The presence of the antibiotic resistance genes, *NEOMYCIN PHOSPHOTRANSFERASE* for the overexpression and *HYGROMYCIN PHOSPHOTRANSFERASE* for CRISPR transformants, was confirmed by PCR. The positive events for overexpression construct were further confirmed by PCR using forward primer for the 35S promoter and reverse primer for *SP3C* gene. Finally, the expression level in *SP3C* overexpressing lines was confirmed by RT-qPCR in RNA extracted from leaves of T_3_ homozygous lines (see below for details). To screen CRISPR/Cas9-generated mutants, the *SP3C* sequence was amplified from DNA extracted of the hygromycin-resistant plants. The PCR products were purified and digested with BsrFI restriction enzyme (cutting site 5′RCCGGY3′), which cuts over the second gRNA sequence in the wild-type genotype, generating two DNA fragments of 605 bp and 347 bp. Transformation events showing only one band would have lost the restriction site and were considered potential mutants. Putative mutants were confirmed by Sanger sequencing in T_3_ homozygous plants. All the primers used are listed in Supplementary Table 2.

To obtain homozygous plants, overexpressing and mutant lines were self-pollinated producing T_1_, T_2_, T_3_ and T_4_ seeds. Progeny tests were carried out on seeds derived from individual plants through foliar spray of antibiotic (400 mg L^-1^ kanamycin for five consecutive days or 100 mg L^-1^ hygromycin for three consecutive days). The progenies (n=30) with 100% antibiotic-resistant plants (assessed visually as absence of leaf yellowing following application) were considered homozygous.

### Real-time quantitative PCR (RT-qPCR)

RNA was purified using ReliaPrep™ RNA Miniprep Systems (Promega), according to the manufacturer recommendations. The cDNA synthesis was performed with SuperScript™ III First-Strand Synthesis System (Invitrogen). Real-Time quantitative PCR reactions (qRT) were performed in a thermocycler Real-Time StepOnePlus PCR (Applied Biosystems) with a final volume of 14 μl using reagent SYBR Green Master Mix (Thermo Fisher Scientific). The melting curves were analyzed for nonspecific amplifications and dimerization of primers. Absolute fluorescence data were analyzed using the software LinRegPCR (Ruijter *et al*., 2009) to obtain the values of quantification cycle (Cq) and calculate the primer efficiency. The abundance of transcripts was normalized against the geometric mean of two reference genes, *TIP4* and *EXPRESSED* and calculated according (Quadrana *et al*., 2013). All the primers used are listed in Supplementary Table 2.

For *SP3C* expression profile the following tissues were used from Micro-Tom (MT) plants: germinated seeds, root tips (≤ 2 cm), hypocotyls, leaf primordia (≤ 0.5 cm), expanding leaf 1 (≤ 2 cm), expanding leaf 2 (≤ 3 cm), mature leaf, flowers (petals + anthers), fruit at immature green 3 stage (IG3) and fruit at red ripe stage (RR).

### Seed germination assay

For germination assays, 72 seeds of each genotype, previously sterilized, were selected and sown in Petri dishes containing 25 mL of MS salts (Murashige and Skoog, 1962) half-strength, vitamins B5, 30 g L^-1^ sucrose and 4 g L^-1^ agar. The pH of the medium was adjusted to 5.7 ± 0.05 and autoclaved for sterilization. The seeds were kept in the dark at 25 ± 1°C throughout the evaluation period. For each genotype four plates with 18 seeds per plate were set and the number of seeds with visible radicle was quantified daily.

### Shoot growth analysis

Flowering time was measured by counting the days from sowing to anthesis and the number of leaves was obtained from the plant base to the first inflorescence. The length and diameter of the internodes, and stem diameter were measured using a mechanical pachymeter (Mitutoyo Vernie^®^, Japan). The leaf angle was determined using a protractor, based on the insertion of the fourth and fifth leaf. The inflorescence morphology was evaluated by counting of flowers number per inflorescence, inflorescence length and number of sepals and petals per flower. At 60 days after germination the plants were harvested and the leaves digitized using a scanner (HP Scanjet G2410) (Hewlett-Packard, Palo Alto, California, USA). Total leaf area was calculated using the software Image-Pro Plus version 4.5 (Media Cybernetics, Silver Spring, USA). Leaves and stems dry weight were obtained after oven drying at 70 °C for 72 h. Specific leaf area (SLA) was calculated through the relationship between leaf area (LA) and leaf dry weight (LDW), as described by the equation: SLA (cm^2^ g^-1^) = LA/LDW. All growth measurements were evaluated in 10 plants per genotype, in MT control, overexpression and knockout lines.

### Root growth analysis

To monitor root development, the plants were grown in vertical PVC cylinders with 30 cm of depth. The cylinders had a hollow bottom covered with a diameter 1.8 × 1.6 mm mesh that allows a quantitative analysis of the root system development and the visualization of root emission over time. The number of secondary roots was quantified daily for thirteen days. The root length, volume and dry weight were evaluated in 55-day-old plants. The root volume was estimated with a graduated cylinder and root dry weight were calculated after oven drying at 70 °C for 72 h.

### Water stress assay

MT plants and either the overexpressing lines (*SP3C*#3) or mutants (*sp3c*#21) were grown paired in single pots. This is a way to ensure the same water supply for both genotypes, as all of them share the same soil water potential during the stress treatment (Bolaños *et al*., 1993). The pots were filled with equal parts of substrate, sand and stones to allow better water drainage and ensure a faster and uniform drying. Watering was suspended after development of two pairs of true leaves in all genotypes, and withheld for eight days, while daily monitoring the water loss of pots. The pots of the well-watered control treatment were irrigated every day to field capacity. For both treatments, well-watered and drought, five replicates were conducted. The seedlings were photographed daily to visually monitor the stress symptoms. The chlorophyll content was estimated throughout the stress period, using a Soil Plant Analysis Development (SPAD).

On the last day of drought, the midday seedling water potential (Ψ) was determined from 12:00 to 14:00, using a Scholander-type pressure chamber (model 1000, PMS Instruments, Albany, NY, USA). Measurements were made in four seedlings per genotype, on drought and control treatments. The relative water content (RWC) also was determined on the eighth day, in four seedlings per genotype. First, the seedlings were weighed to obtain fresh weight (FW) and then submerged in distilled water for 24 h. After this time, the seedlings were weighed again to obtain turgid weight (TW) and dried at 70°C for 48 h to determine dry weight (DW). The RWC was determined by the equation: RWC=[(FW–DW)/(TW–DW)] × 100. After this treatment, the remaining seedlings were rehydrated and the percentage of recovered seedlings evaluated. Total leaf area, specific leaf area and dry weight were evaluated in all surviving seedlings.

### Statistical analysis

The experimental design was completely randomized. Data were submitted to analysis of variance (ANOVA) and the means were compared by Tukey test at 5% level of significance (*P* ≤ 0.05) using Sisvar^®^ software (version 5.6).

## Results

### Genomic characterization of the CETS gene family in tomato

The phylogenetic grouping of the CETS proteins in tomato and *Arabidopsis* shows that the *CETS* genes are distributed into three well-defined clades: *FT*-like, *TFL1*-like and *MFT*-like, based on their sequence similarity to the founding members in *Arabidopsis*. *TFL*1-like groups the flowering repressors and *FT*-like includes the flowering inducers (Fig. 1 and Supplementary Fig. S1). Sequence comparison between *CETS* genes of the wild ancestor of tomato *Solanum pimpinellifolium* and domesticated tomato (*S. lycopersicum*) revealed a high structural conservation between paralogues in both species (Fig. 1). All of them have four exons and three introns, and the main difference between paralogues within each species is the variation in intron length (Fig. 1). Previous work uncovered those polymorphisms leading to gene loss of function in some members of the family underlie important domestication and improvement traits. For instance, determinate growth habit is produced by a single nucleotide change in *SP* (Pnueli *et al*., 1998), whereas deletions in the *cis*-regulatory region of *SP5G* and the fourth exon of *SP11B.1* synergistically reduce the short-day flowering response leading to day-neutral flowering (Song *et al*., 2020) (Fig. 1).

**Fig. 1.**
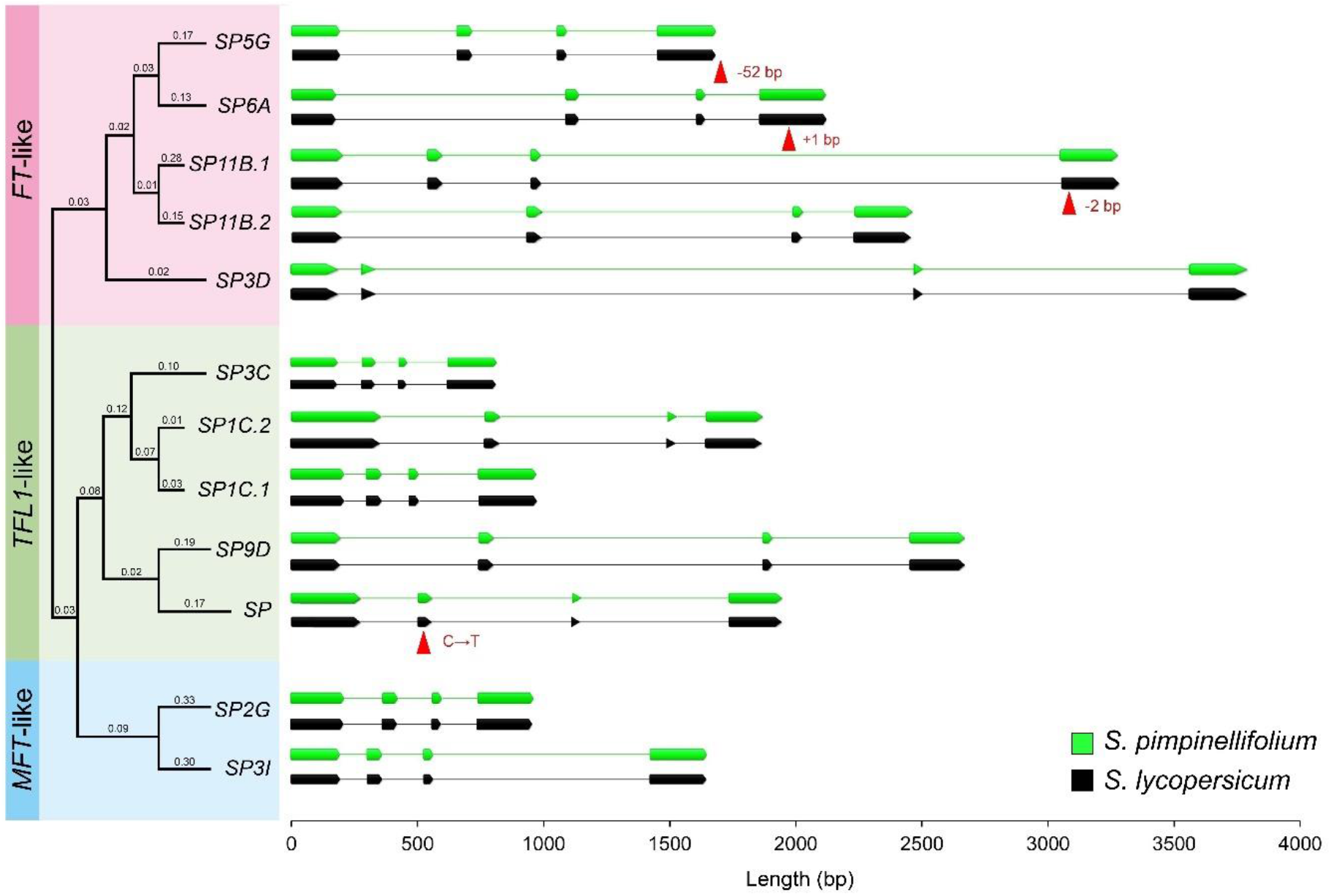
Allelic variation in the *CETS* gene family underpins tomato domestication and improvement. Left: Phylogenetic reconstruction obtained from alignment of tomato *CETS* (*CENTRORADIALIS/TERMINAL FLOWER 1/SELF-PRUNING*) gene sequences. The three clades, based on the *Arabidopsis thaliana* homologues, are: *MOTHER OF FLOWERING LOCUS T* (*MFT*)-like genes (blue), *TERMINAL FLOWER 1* (*TFL1*)-like genes (green) and *FLOWERING LOCUS T* (*FT*)-like genes (pink). Values show the mean substitution rate per nucleotide. Right: Schematic alignment comparing the *CETS* genes structure in the putative tomato ancestral species *Solanum pimpinellifolium* (green) and domesticated tomato *Solanum lycopersicum* cv. Heinz 1706 (black). Diagrams show exons (boxes) and introns (lines). The previously described polymorphisms relevant to tomato domestication and improvement are indicated by red arrows (see text for details). The full sequences are listed in Supplementary Table S1.

We previously showed that *SP3D* (*SFT*) coordinates flowering with stomatal conductance and water loss (Robledo et al, 2020), thus, here we decided to explore a potential role for other members of the *CETS* family in plant responses to drought.

The wild tomato relative *S. pennellii* is endemic of semi-arid and arid regions of the Andes Mountains and is therefore highly drought-tolerant (Taylor, 1986). We first analysed the effect of progressive water deficit in the soil on the transcription of *CETS* genes in leaves of *S. pennellii* and *S. lycopersicum* (Fig. 2A). We found that *SP2G, SP3C* responded to the drought treatment by increasing their expression in both species. Interestingly, *SP11B.1* showed a decrease in *S. pennellii* in the drought treatment, but no expression in domesticated tomato, as previously described (Song *et al*. 2020). *SP3C* showed the highest expression increase during drought, so to gain further insight into its involvement in drought responses, we analyzed the promoter region of the *SP3C* gene, and compared it with that of the two main flowering regulators, *SP* and *SP3D (SFT)*. Some motifs were common between two or three of these genes such as anaerobic induced motifs, light response and hormonal responses. Interestingly, MYB drought-responsive and seed-specific motifs were found exclusively in the *SP3C* promoter (Fig. 2B). The responsiveness to drought and the presence of the seed and drought responsive motifs in its promoter led us to produce a more in-depth functional characterization of *SP3C*.

**Fig. 2.**
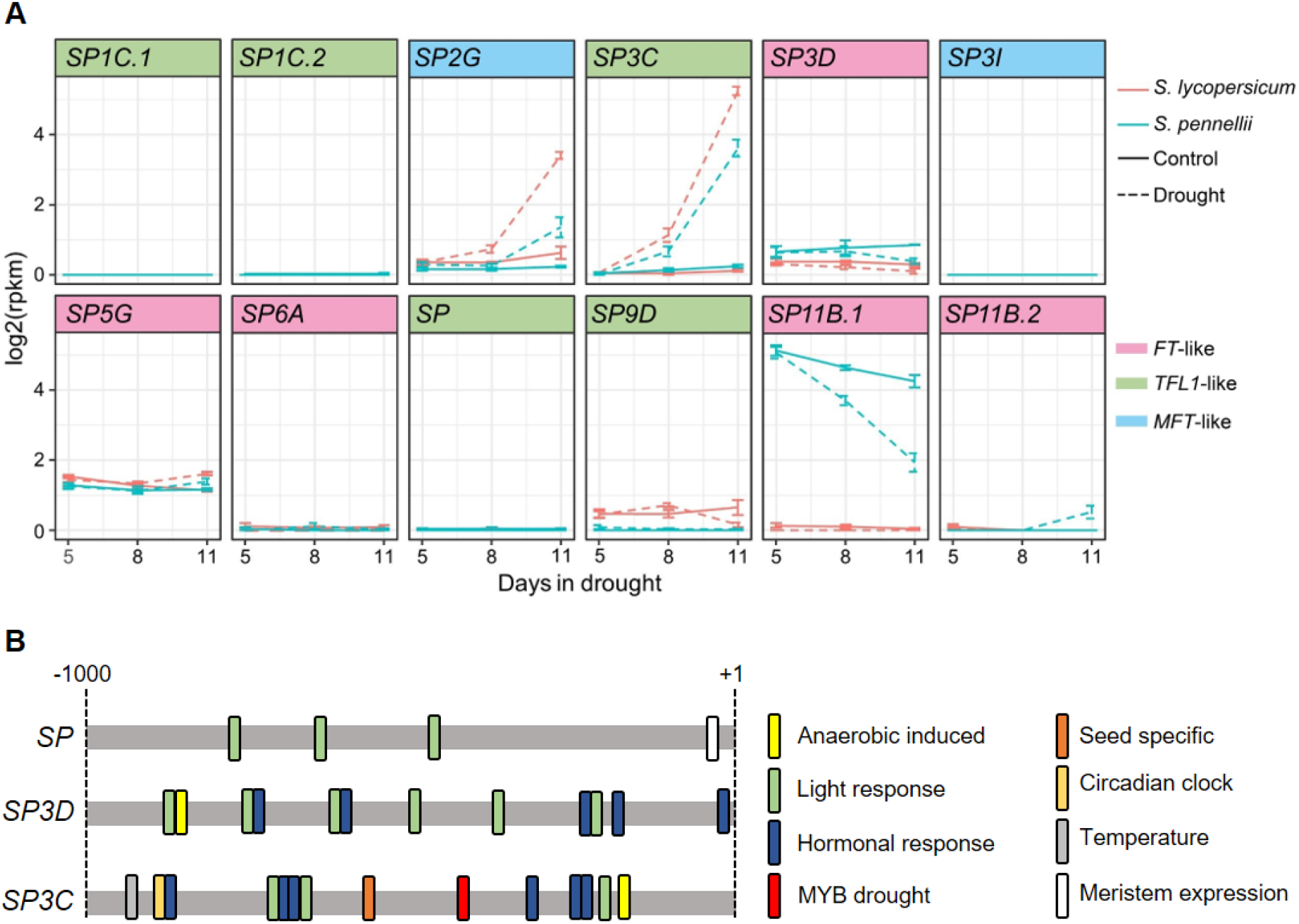
*SP3C* expression is induced by drought in tomato. (A) Relative expression of *CETS* genes in leaves of *S. pennellii* (turquoise) and *S. lycopersicum* (magenta), collected at three points after suspension of watering (drought) compared to well-watered (control) plants. *SP2G* and *SP3C* were upregulated in both species along the time-course of drought, *SP3C* showing a more expressive response, and *SP11B.1* was downregulated as the drought progressed, but only in wild specie *S. pennellii*. (B) Putative transcription factor binding sites found in the promoter region (1000 bp upstream of the start codon) of the *SP*, *SP3D* (*SFT*) and *SP3C* genes from *S. lycopersicum*. Anaerobic induced, light response and hormonal response motifs are shared in two or three of these genes. While, MYB drought responsive and seed-specific motifs were exclusively found in the *SP3C* promoter.

### SP3C characterization and generation of overexpressing and mutant lines

*SP3C* encodes a putative PEBP-like protein of 175 amino acids and belongs to the *TFL1-* like clade (Fig. 1), suggesting a potential role as flowering repressor. This putative role is reinforced by the presence of a repressor amino acid signature, a histidine residue homologous to the *AtTFL1* His-88 (Supplementary Fig. S1) (Hanzawa *et al*., 2005; Hedman *et al*., 2009). We analyzed the expression levels of *SP3C* mRNA in different organs and developmental stages of tomato cv. Micro-Tom (MT). The most prominent levels of expression were found in ‘Immature Green 3’ stage fruits (IG3) and roots (Fig. 3A). We next generated overexpressing transgenic and loss-of-function mutant lines. Two independent transformation events, *35S*::*SP3C*#1 and *35S::SP3C*#3, hereafter named *SP3C*#1 and *SP3C*#3, showed *SP3C* expression levels between 700- and 800-fold higher than the MT control in the T_3_ homozygous generation (Fig. 3B). For the loss-of-function experiment, three independent CRISPR-mediated *SP3C* homozygous events were obtained, *sp3c*#12, *sp3c*#21 and *sp3c*#24, and the mutations were confirmed by Sanger sequencing (Fig. 3C). At the first gRNA site (Target 1) on the first exon, all three mutants had a single nucleotide insertion (+G) 172 bp downstream of the start codon, that resulted in disruption of the reading frame. At the site targeted by the second gRNA (Target 2), the line *sp3c*#12 had a 3-bp deletion (CCG), whereas lines *sp3c*#21 and *sp3c*#24 showed a 55-bp deletion (Fig. 3C). These mutant alleles resulted in a premature stop codon and a predicted protein of 74 amino acids (Supplementary Fig. S2). With these genotypes, we conducted a broad phenotypic characterization and investigated the effects of *SP3C* in phenology, growth and morphology of tomato plants (Fig. 3D).

**Fig. 3.**
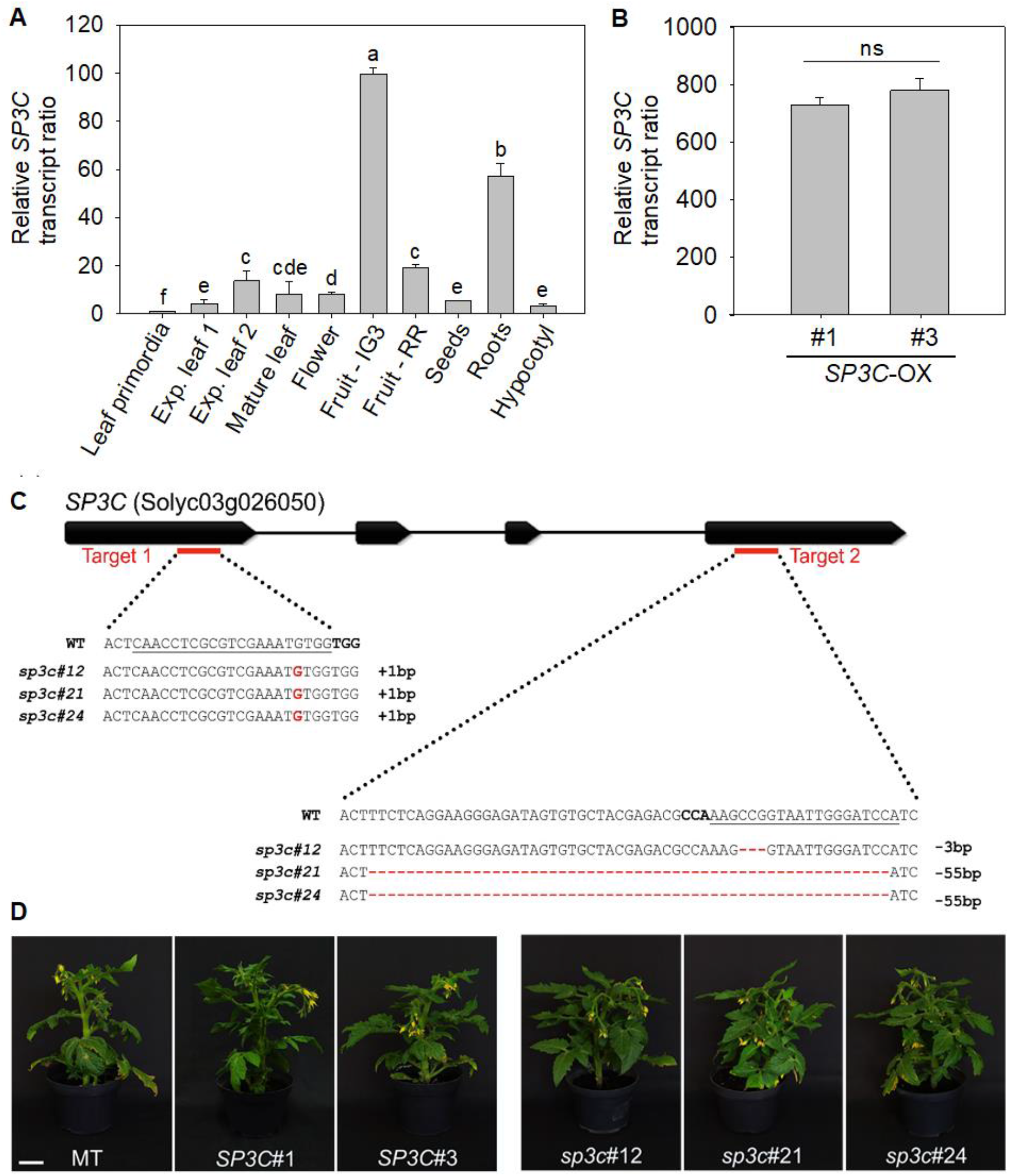
Generation of new allelic variants for *SP3C* in tomato cv. Micro-Tom (MT). (A) Expression profile of *SP3C* in leaf primordia, expanding leaf 1, expanding leaf 2, mature leaf, flower, fruit at immature green 3 stage (IG3), fruit at red ripe stage (RR), seeds, roots and hypocotyl. Data were normalized against the leaf primordia sample. Values are means ± SE (n=3) and different letters indicate significant differences between each analysed sample (*P*<0.05). (B) Relative *SP3C* transcript levels in young leaves of *35S*::*SP3C*#1 (*SP3C*#1) and *35S*::*SP3C*#3 (*SP3C*#3) overexpressing lines. Data were normalized against the MT wild-type. Values are means ± SE (n=3). (C) CRISPR/Cas targeting to the first and fourth exon of *SP3C* using two guide RNAs (gRNAs); target 1 and target 2, red lines. The sequences below show the *sp3c*#12, *sp3c*#21 and *sp3c*#24 alleles identified from three independent transgenic plants, T_3_ generation. gRNAs sequences are shown underlined, protospacer-adjacent motifs (PAMs) are highlighted in bold, insertions in red and deletions in red dashes. The sequence length differences compared to MT are shown to the right (D) Representative plants of MT, *SP3C* overexpressing lines (*SP3C*#1; *SP3C*#3) and *sp3c* loss-of-function mutant lines (*sp3c*#12; *sp3c*#21; *sp3c*#24) with 60 days after sowing. Scale bars, 5 cm.

### SP3C is a repressor of seed germination

While screening the overexpression lines for homozygous insertions we noticed a reduction in seed germination rate, especially in the *SP3C* overexpression line *SP3C*#1 (Supplementary Fig. S3). These observations, together with the presence of seed-specific expression motifs in the *SP3C* promoter (Fig. 2B), led us to quantify seed germination rate in a controlled assay. Homozygous *SP3C* overexpressing lines displayed a significant delay in germination compared to wild-type MT plants (Fig. 4A). After 12 days, over 96% of the MT seeds and mutants germinated; however only 21% and 78 % of *SP3C*#1 and *SP3C*#3, respectively, showed radicle protrusion through the seed testa. Moreover, a slight acceleration in mutant seeds germination between the 4th and the 5th day after sowing was observed as well for the loss-of-function lines *sp3c*#12 and *sp3c*#21. Our results indicate *SP3C* repress seed germination.

**Fig. 4.**
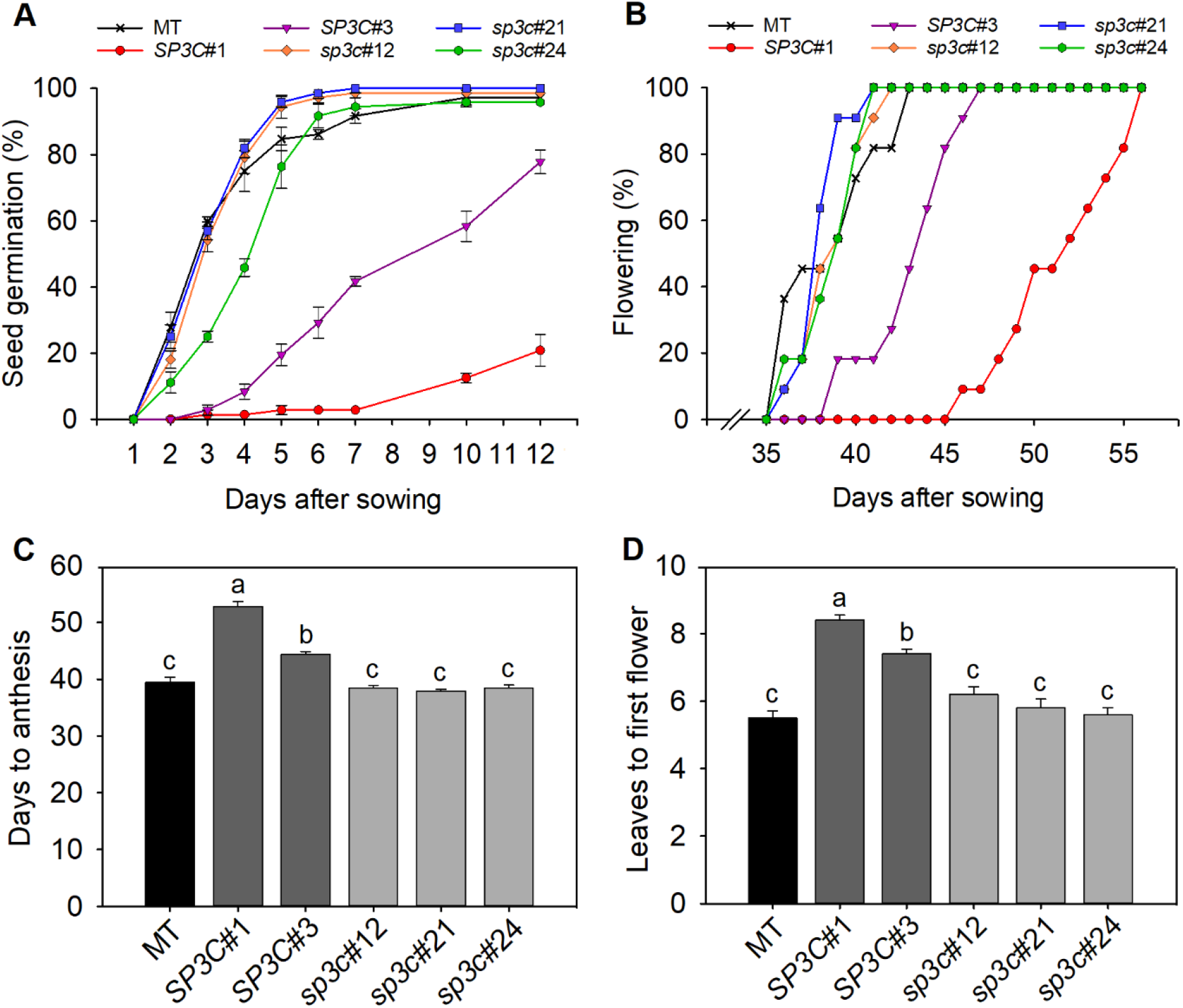
*SP3C* represses seed germination and delays flowering. (A) Germination time in tomato cultivar Micro-Tom (MT), *35S*::*SP3C*#1 (*SP3C*#1) and *35S::SP3C#3 (SP3C#3*) overexpression lines and *sp3c*#12, *sp3c*#21 and *sp3c*#24 mutant lines. The number of germinated seeds (in percentage) was evaluated daily for 12 days. Data are means ± SE of four biological replicates with 18 seeds each. (B) Flowering time in MT, overexpression lines and mutants. (C) Mean number of days from sowing to anthesis. Data are means ± SE (n = 10 plants). Different letters indicate statistically significant differences (ANOVA + Tukey’s test, P <0.05). (D) Number of leaves to first inflorescence. Data are means ± SE (n = 12 plants). Different letters indicate statistically significant differences (ANOVA + Tukey’s test, P<0.05).

### SP3C is a flowering repressor

Most vegetative growth parameters showed no evident alterations in both transgenic *35S::SP3C* and *sp3c* mutant lines compared to the MT control (Supplementary Table S3 and Fig. S4). However, *SP3C*-overexpressing plants showed delayed flowering and a greater number of days to anthesis, whereas *sp3c* mutants had a subtle but consistent acceleration of flowering (Fig. 4B, C). The number of leaves to the first inflorescence was higher in *35S::SP3C* lines, while it remained unchanged in *sp3c* mutants (Fig. 4D). These results suggest that *SP3C*, similarly to other members of the *CETS* family, contribute to flowering control; particularly, acting as a floral repressor in tomato.

In addition, we also analysed parameters related to inflorescence morphology and observed that although the floral structure was not altered, the inflorescence morphology was modified in the overexpressing lines. The *35S::SP3C* lines had a greater number of flowers per inflorescence unit and a longer inflorescence (Supplementary Table S3 and Fig. S5).

### SP3C reduces root length and stimulates root branching

The presence of drought-responsive motifs in the *SP3C* promoter and the high expression level in roots hinted at a putative role of *SP3C* in root development. To better explore this, we conducted an experiment under controlled conditions to assess root growth (Fig. 5A). We observed that one of the *35S::SP3C* overexpressing events and two *sp3c* mutants showed reduced and increased root length in relation to MT control, respectively (Fig. 5B, C). A time-course of secondary root development revealed that *SP3C*#3 plants had faster root branching than MT, whereas *SP3C*#1 lagged behind MT, with all three genotypes eventually reaching ~100 secondary roots 32 days after transplanting. On the other hand, secondary root development was severely compromised in the mutant lines: all of them showed a slower rate of root branching and at the end of the evaluation period *sp3c*#12 had 43% (*p* = 0.1236), *sp3c*#21, 58% (*p* = 0.0028), and *sp3c*#24, 65% (*p* = 0.0004) lower number of secondary roots than MT (Fig. 5D). Root volume and dry mass were also significantly reduced in the mutants (Supplementary Fig. S6). Thus, these data revealed that *SP3C* represses principal root growth and increases root ramification.

**Fig. 5.**
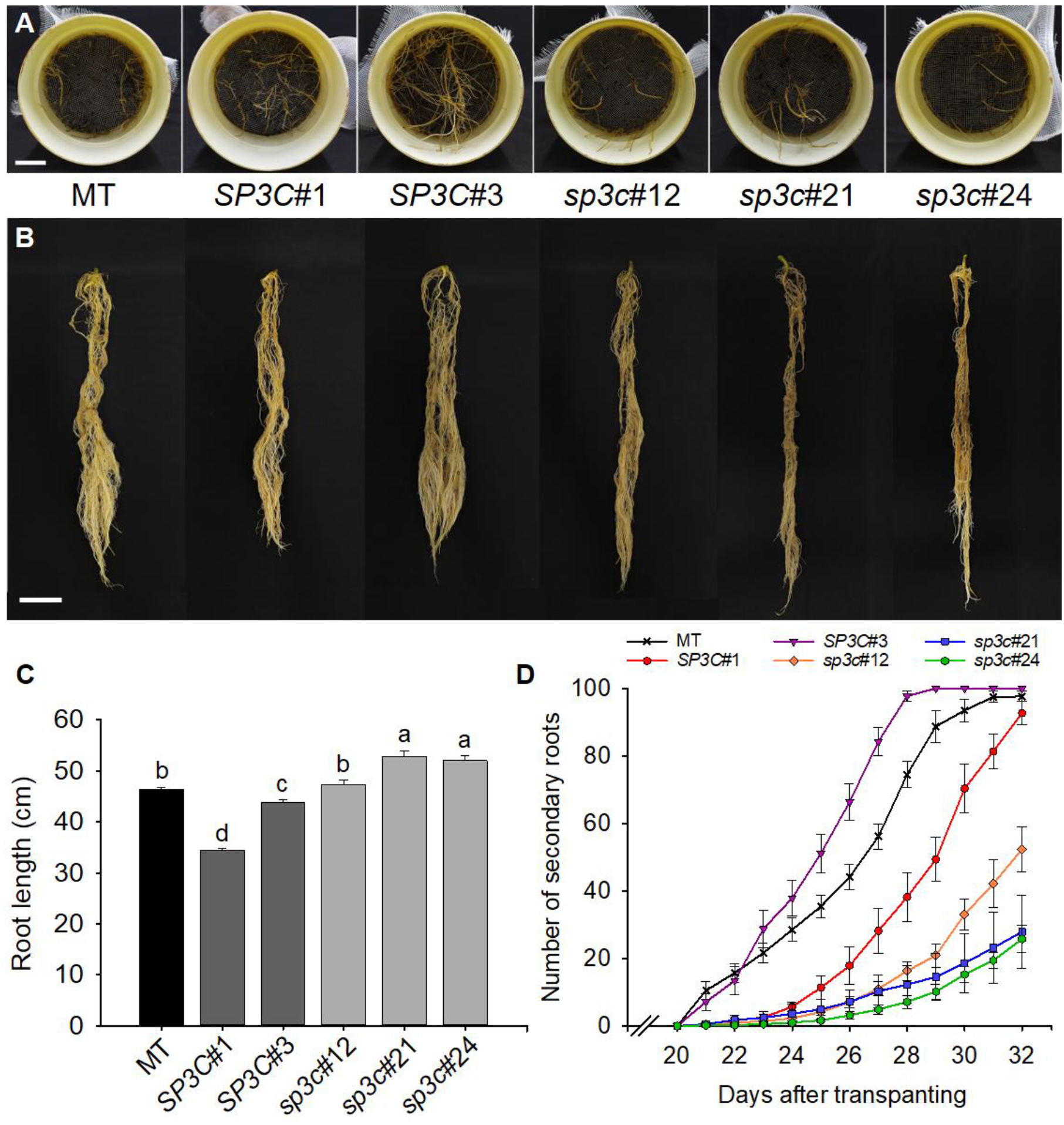
*SP3C* regulates root architecture. (A) Representative photo of secondary roots emission in tomato cultivar Micro-Tom (MT), *SP3C* overexpressing lines (*SP3C*#1; *SP3C*#3) and *sp3c* loss-of-function mutant lines (*sp3c*#12; *sp3c*#21 and *sp3c*#24) 32 days after transplanting for vertical PVC cylinders. Scale bars, 2 cm. (B) Root morphology at 50 days after transplanting (dat). Scale bars, 2 cm. (C) Root length measurements at 50 dat. Data are means ± SE (n = 10 plants). Different letters indicate statistically significant differences (ANOVA + Tukey’s test, P <0.05). (D) Time course of secondary root emission. The number of visible roots was measured at 15 dat for a period of 13 days.

### SP3C improves the drought response in seedlings

We showed that *SP3C* expression is increased under drought conditions, which, coupled to its impact on seed germination and root growth, could indicate a possible involvement in plant responses to water stress. To further investigate the relation between *SP3C* and drought, we conducted a water stress assay in 22-day-old seedlings of two selected lines: one overexpression (*SP3C*#3) and one mutant (*sp3c*#21), for eight days. The soil water loss was monitored daily during water stress. On the eighth day, the drought treatment pots had a loss of approximately 80% in their water content compared to the start of treatment (Supplementary Fig. S7).

During the time-course of progressive soil drying, *sp3c*#21 mutant seedlings showed more intense leaf wilting symptoms, while *SP3C*#3 visually appeared to be less affected by drought (Fig. 6A). On the seventh day of water stress, *sp3c*#21 mutant seedlings were fully wilted, considerably more than MT control plants, and *SP3C*#3 seedlings began to show noticeable symptoms of leaf rolling (Fig. 6A). We ensured that all plants from the drought treatment were under stress by measuring water potential (Ψ), relative water content (RWC) and chlorophyll content (Fig. 6B-E). The Ψ and RWC of genotypes under drought was consistently lower than control plants.

**Fig. 6.**
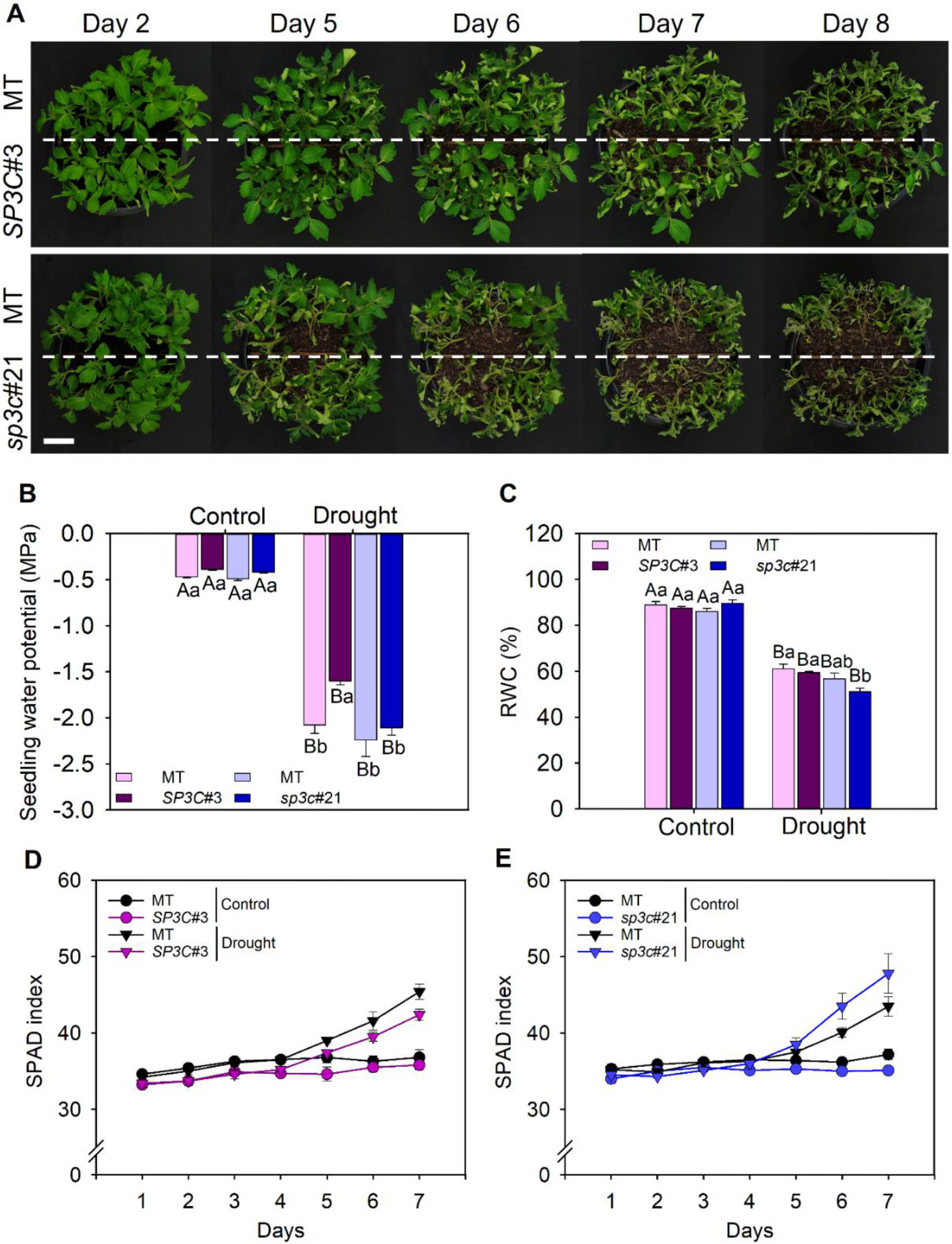
*SP3C* improves tomato seedlings response during the drought. (A) Representative photo of genotypes Micro-Tom (MT) grown together with overexpressing line (MT/*SP3C*#3) and Micro-Tom (MT) grown together with mutant line (MT/*sp3c*#21) throughout the assay. The leaf wilt symptoms were recorded daily, during eight days of drought. Visually, the seedlings began to show symptoms of water deficit on 5^th^ day and were progressively intensified until 8^th^ day of drought. The overexpression line *SP3C*#3 showed more subtle symptoms of winding and leaf wilt, while in mutant *sp3c*#21 the symptoms were clearly more intense. Scale bars, 2 cm. (B) Seedling water potential measured at 12:00 on the 8^th^ day after suspension of watering, in MT/*SP3C*#3 (purple) or MT/*sp3c*#21 (blue). (C) Relative water content (RWC) obtained by the relation between fresh, turgid and dry weight of seedlings, on the 8^th^ day after stress. Data are means ± SE n = 5 plants. Different letters indicate statistically significant differences (ANOVA + Tukey’s test, P <0.05). (D) Chlorophyll content estimated by SPAD along the drought in MT/*SP3C*#3 and (E) MT/*sp3c*#21.

Our results suggest that the greater tolerance of *SP3C*#3 line to drought was associated with higher Ψ and higher water status at the end of the drought treatment. We also showed that the tolerance of this line is exclusively related to the increase in *SP3C* expression and not to other factors that could interfere in the genotype-specific response. Thus, we ensured that there were no discrepancies in total leaf area, specific leaf area and biomass between *SP3C*#3 lines and *sp3c*#21 mutants, which could impact the stress response (Supplementary Fig. S7).

## Discussion

The transition from vegetative to reproductive growth is a key determinant of plant architecture and agronomic performance, as it conditions photosynthetic source-sink relations and productivity. Flowering time has therefore been one of the main breeding targets, allowing crops to perform better under specific environmental conditions and driving the expansion of agricultural areas to a wider range of latitudes and climates (Chen *et al*., 2020). It had long been known that a flowering stimulus is produced in the leaves and translocated to the apical meristem, where it triggers the switch from vegetative to reproductive growth (Zeevaart, 2006). This signal, the ‘florigen’, was elusive for many decades but eventually shown to be small polypeptides (<200 amino acids) encoded by the *CETS (CENTRORADIALIS, TERMINAL FLOWER 1, SELF PRUNING*) gene family (Susila *et al*., 2021). *CETS* family has undergone expansion via duplication and subfunctionalization in angiosperms (Wickland and Hanzawa, 2015). In spite of the partial functional redundancy of the orthologs within each sub-clade, in some species it has been shown that each one fulfils specific roles to fine-tune plant development in response to specific environmental cues.

Tomato is a valuable model to gain further functional insight into the *CETS* gene family (Carmel-Goren *et al*., 2003). A large-scale genomic analysis demonstrated that three of the twelve *CETS* genes (*SP3C, SP5G* and *SP6A*) are located within genomic regions comprising putative domestication sweeps and six others (*SP1C.1, SP1C.2, SP3D, SP, SP11B.1* and *SP11B.2*) in putative improvement sweeps (Lin *et al*., 2014). This suggests that selection of allelic variation on genes of the *CETS* family has been a key driver of the changes operated along the transitions between *S. pimpinellifolium* and *S. lycopersicum* var. *cerasiforme* (domestication), and between *S. lycopersicum* var. *cerasiforme* and modern fresh fruit and processing varieties (improvement) of *S. lycopersicum* (Razifard *et al*., 2020). A single amino acid change in the *SP* gene (Fig. 1) leads to determinate growth, which includes alteration in the auxin signaling (Silva *et al*., 2018), and underpins the development of processing tomato varieties that are suitable for mechanical harvest (Pnueli *et al*., 1998). In their natural habitats wild tomatoes respond to photoperiodic cues and show a strong short-day response to flowering (Taylor, 1986). Mutations in the *SP5G* and *SP11B.1* loci (Fig. 1) were selected to suppress the photoperiodic response and are found in almost completely day neutral tomato varieties grown in higher latitudes (Soyk *et al*., 2017; Zhang *et al*., 2018; Song *et al*., 2020). Lastly, a loss-of-function mutation in the fourth exon of *SP6A* is ubiquitous in domesticated tomatoes (Fig. 1). Its functional significance is hitherto unknown, but *SP6A*, in conjunction with *SP5G*, is the main controller of tuberization in response to day length in potatoes (Abelenda *et al*., 2016).

A deeper understanding of the expression patterns and phenotypic effects of the genes of the *CETS* family would provide potentially useful targets for breeding using the state-of-the-art gene-editing suite of tools (Gasparini *et al*., 2021). A loss-of-function mutation in *SFT (SP3D*) improves the vegetative-to-reproductive growth balance when in heterozygosis and combined with the *SP* loss-of-function allele (*sp*) in homozygosis (Krieger *et al*., 2010). Likewise, the tomato plant architecture can be fine-tuned and optimized by targeted mutations in the promoter region of the *SP* gene (Rodríguez-Leal *et al*., 2017). This suggests that considerable room for improvement of plant growth and agronomic performance exists by manipulation of *CETS* family genes, not only in tomato but also in other crop species (McGarry and Ayre, 2012).

Here, we first showed that the expression of the *SP2G* and *SP3C*, orthologs of *Arabidopsis MFT* and *TFL1*, is strongly induced in the leaves of the drought-resistant wild species *S. pennellii* and domesticated tomato upon suspension of watering (Fig. 2A). The increased expression pattern in proportion to the time under drought was unique to both these genes and not observed in the other paralogues of the *CETS* family. We previously showed that the *SFT* (*SP3D*) is responsible for controlling water-use efficiency in tomato by altering stomatal responses upon flowering (Robledo *et al*., 2020). This indicates that the *CETS* family genes have pleiotropic roles that go beyond the control of plant development and may include modulation of water relations.

We focused our analyses on dissecting the function of the *SP3C* gene, and demonstrated that it pleiotropically controls seed germination, flowering time and root development. Increased *SP3C* expression in transgenic lines led to delayed in seed germination and flowering. On the other hand, using CRISPR/Cas9 to create targeted mutations in the coding region of the *SP3C* gene accelerated seed germination and subtly shortened flowering time. Seed germination time is a fundamental domestication trait, as most wild species have genetic and physiological mechanisms to induce strong dormancy (Chahtane *et al*., 2017; Penfield, 2017). Correct timing of germination to coincide with favorable environmental conditions can result in increased survival and fitness (Wang *et al*., 2018). However, extended seed dormancy is not a desirable trait for crops, as it can lead to erratic and unpredictable germination rates (Rodríguez *et al*., 2015), preclude the better use of fertilizers and extend the growth cycle. *MFT* represses seed germination under far-red light by modulating ABA and hormone responses (Xi *et al*., 2010; Vaistij *et al*., 2018). Recent work in *Arabidopsis* has shown that ectopic *FT* expression in seeds increased dormancy by modulating gibberellin biosynthesis (Chen *et al*., 2021). Further studies would clarify whether the modulation of seed germination by *SP3C* could involve hormonal alterations.

The transition to flowering in angiosperms is controlled through a mechanism that integrates environmental and endogenous signals that converge on the expression of *CETS* genes (Cao *et al*., 2021). *CETS* genes encode flowering repressors (similar to *TFL1*) and flowering inducers (similar to *FT*), which can differ between them by as little as a single amino acid (Hanzawa *et al*., 2005), and compete for the binding site of *FLOWERING LOCUS D (FD)*, a bZIP transcription factor with localized expression in the meristem (Collani *et al*., 2019). Binding of FD to a CETS protein and adaptor proteins of the 14-3-3 family leads to formation of the Flowering Activator Complex (FAC) (Ho and Weigel, 2014), which drives the reprogramming of the meristem and its conversion from vegetative to floral (Périlleux *et al*., 2019). Here, we showed that *SP3C* is a negative regulator of flowering, which is consistent with its placing in the *TFL1*-like clade of proteins that in tomato includes the anti-florigenic signals *SP* and *SP9D*. Moreover, the high expression of *SP3C* in roots and immature fruits immediately points to functions beyond the control of flowering time.

Root growth patterns were also affected by alteration of *SP3C* function. High *SP3C* expression led to reduced root length but increased root branching and secondary roots. Loss of *SP3C* function had the opposite effect of increasing root length but decreasing secondary root formation. Previous work showed that *TFL1* genes are expressed in the roots of many species, including *Arabidopsis*, where they can act as repressors of root growth, leading to reduced root length (Bouché *et al*., 2016). The expression of *CETS* family genes in roots suggests a potential mechanism whereby the coordination of flowering time and root growth is achieved. In many species (*e.g* sugar beet and potato) the transition to flowering triggers a halt in root growth that is fundamental to coordinate resource allocation and guarantee reproductive success (Pin *et al*., 2010; Navarro *et al*., 2011; Abelenda *et al*., 2016). Further investigation of the specific roles of diverse *CETS* paralogues on this phenomenon could provide a novel target to create root systems that are tailored to specific environments based on the design of ideotypes and their creation using gene editing tools (Lynch, 2019).

The pleiotropic effects described for many members of the *CETS* family, on which we have expanded here, would make them important drivers of domestication and breeding, as relatively simple genetic changes could produce multiple beneficial traits simultaneously. The discovery and characterization of a flowering repressor with pleiotropic effects on seed germination and root growth represents an advance in the effort to link genotypes to agronomically relevant traits in horticultural crops. Further work is required to shed more light on the effect of drought on *SP3C* expression and function. In summary, our study provides a new target for simultaneous modulation of seed germination, flowering and root growth that could be fine-tuned for agronomic gain.

## Supplementary data

The following supplementary data are available at *JXB* online.

*Table S1*. Sequences used for phylogenetic analysis.

*Table S2*. Oligonucleotides DNA sequence for PCR primers used in this study.

*Table S3*. Growth parameters measured in Micro-Tom (MT), overexpression lines and mutants.

*Fig. S1*. Characterization of the *SP3C* gene.

*Fig. S2*. Amino acid sequences of the *sp3c* mutant alleles.

*Fig. S3. SP3C* acts as a repressor of seed germination.

*Fig. S4*. Tomato plants harboring allelic variations in the *SP3C* gene.

*Fig. S5*. High levels of *SP3C* increase the flowers number and the inflorescence length.

*Fig. S6*. Impacts of *SP3C* on root grown and development.

*Fig. S7. SP3C* improves tomato seedling response to drought.

## Acknowledgements

We thank the support and contributions of the UFV Plant Physiology Graduate Program.

## Author contributions

L. E.P.P., M.R. and A.Z. designed study. J.d.R.M., A.Q., B.S.L., J.M.R., S.J.C., M.H.V. and M. Ry. performed experiments. M.Ry. and J.M.J-G. analyzed data. D.M.R., J.M.G-G., L.E.P.P., M.R. and A.Z. acquired funding and managed research personnel. J.d.R.M. and A.Z. wrote the manuscript. All authors commented, edited and approved the final manuscript.

## Conflict of interest

The authors declare no conflict of interest.

## Funding

This work was funded in part by a grant (RED-00053-16) from the Foundation for Research Assistance of the Minas Gerais State (FAPEMIG, Brazil) and by funding from the German Research Foundation under the German-Israeli Project Cooperation program (DFG DIP project number FE552/12-1 awarded to J.M.J.-G.).

### Disclaimer

The views and opinions expressed in this work do not necessarily reflect the official policy or position of the authors’ respective employer or government.”. Mention of any trade name, proprietary product or specific equipment does not constitute a guarantee or warranty by USDA-ARS and does not imply its approval to the exclusion of other products that may also be suitable. The USDA-ARS is an equal opportunity and affirmative action employer and all agency services are available without discrimination.

## Data availability

All data supporting the findings of this study are available within the paper and within its supplementary data published online. No restrictions are placed on the materials, such as material transfer agreements.

## Notes

### Competing Interest Statement

The authors have declared no competing interest.

