## Supplementary materials for "*SELF PRUNING 3C* is a flowering repressor that modulates seed germination, root architecture and drought responses"

[1]Departamento de Biologia Vegetal, Universidade Federal de Viçosa, Viçosa, Brazil.

[2]Departamento de Botânica, Universidade de São Paulo, São Paulo, Brazil.

[3]United States Department of Agriculture, Plant Science Research Unit, St Paul, MN 55108, USA

[4]Department of Agronomy and Plant Genetics, University of Minnesota, St. Paul, MN 55108, USA

[5]Center for Plant Precision Genomics, University of Minnesota, St. Paul, MN 55108, USA

[6]Center for Genome Engineering, University of Minnesota, St. Paul, MN 55108, USA

[7]Departamento de Ciências Biológicas, Escola Superior de Agricultura "Luiz de Queiroz", Universidade de São Paulo, Piracicaba, Brazil.

[8]Max Planck Institute for Plant Breeding Research, Cologne, Germany

[9]Centro de Biotecnologia y Genomica de Plantas (CBGP), Madrid, Spain

‡Corresponding author:

Zsögön, A.

**Supplementary Table S1.** Sequences used for phylogenetic analysis.

| Locus | Species | Protein name | Group |
|---|---|---|---|
| fgenesh1_pg.9_152 | *Coccomyxa subellipsoidea* C-169 | - | Chlorophyta |
| estExt_fgenesh1_pg.C_20153 | *Coccomyxa subellipsoidea* C-169 | - | Chlorophyta |
| AT1G18100 | *Arabidopsis thaliana* | *At*MFT | MFT-like |
| AT1G65480 | *Arabidopsis thaliana* | *At*FT | FT-like |
| AT2G27550 | *Arabidopsis thaliana* | *At*ATC | TFL1-like |
| AT4G20370 | *Arabidopsis thaliana* | *At*TSF | FT-like |
| AT5G03840 | *Arabidopsis thaliana* | *At*TFL1 | TFL1-like |
| AT5G62040 | *Arabidopsis thaliana* | *At*BFT | TFL1-like |
| Dusal.0044s00009 | *Dunaliella salina* | - | Chlorophyta |
| Solyc01g009560 | *Solanum lycopersicum* | *Sl*SP1Ca | TFL1-like |
| Solyc01g009580 | *Solanum lycopersicum* | *Sl*SP1Cb | TFL1-like |
| Solyc02g079290 | *Solanum lycopersicum* | *Sl*SP2G | MFT-like |
| Solyc03g026050 | *Solanum lycopersicum* | *Sl*SP3C | TFL1-like |
| Solyc03g063100 | *Solanum lycopersicum* | *Sl*SP3D/*Sl*SFT | FT-like |
| Solyc03g119100 | *Solanum lycopersicum* | *Sl*SP3I | MFT-like |
| Solyc05g053850 | *Solanum lycopersicum* | *Sl*SP5G | FT-like |
| Solyc05g055660 | *Solanum lycopersicum* | *Sl*SP6A | FT-like |
| Solyc06g074350 | *Solanum lycopersicum* | *Sl*SP | TFL1-like |
| Solyc09g009560 | *Solanum lycopersicum* | *Sl*SP9D | TFL1-like |
| Solyc11g008640 | *Solanum lycopersicum* | *Sl*SP11Ba | FT-like |
| Solyc11g008650 | *Solanum lycopersicum* | *Sl*SP11Bb | FT-like |
| Solyc11g008660 | *Solanum lycopersicum* | *Sl*SP11Bc | FT-like |

MFT-like: *MOTHER OF FT AND TFL1*, FT-like: *FLOWERING LOCUS T*, TFL1-like: *TERMINAL FLOWER1*.

**Supplementary Table S2.** Oligonucleotides DNA sequence for PCR primers used in this study. The sequence of oligonucleotides used for cloning, genotyping and RT-qPCR are listed.

| Target gene | Forward sequence (5'-3') | Reverse sequence (5'-3') | Purpose |
|---|---|---|---|
| *SP3C* | ATGTCTTCTAGAAGTACTTG | CAAACACCATATGTATATAT | cloning/ genotyping |
| *M13* | GTAAAACGACGGCCAG | CAGGAAACAGCTATGAC | cloning |
| *pCaMV 35S* | CCTCGGATTCCATTGCCCA | ---- | genotyping |
| *NPTII* | GAGGCTATTCGGCTATGACTGG | ATCGGGAGCGGCGATACCGTA | genotyping |
| *HPT* | CTATCGGCGAGTACTTCTAC | GATCCCCATGTGTATCACTG | genotyping |
| *SP3C* | CATGAGATCTGCTTATACACTC | CATAACGATGGATCCCAATTAC | qPCR |
| *TIP4* | GCTGCGTTTCTGGCTTAGG | ATGGAGTTTTTGAGTCTTCTGC | qPCR |
| *EXP* | GCTAAGAACGCTGGACCTAATG | TGGGTGTGCCTTTCTGAATG | qPCR |

**Supplementary Table S3.** Growth parameters measured in Micro-Tom (MT), overexpression lines (*SP3C* #1; *SP3C* #3) and mutants (*sp3c* #12; *sp3c* #21 and *sp3c* #24).

| **Parameters** | **MT** | ***SP3C* #1** | ***SP3C* #3** | ***sp3c* #12** | ***sp3c* #21** | ***sp3c* #24** |
|---|---|---|---|---|---|---|
| Plant height (cm) | 11.51 ± 0.42b | 12.03 ± 0.33ab | 12.70 ± 0.28a | 12.88 ± 0.19a | 11.78 ± 0.25ab | 11.42 ± 0.22b |
| **Internode length (cm)** | | | | | | |
| Internode 4 | 1.29 ± 0.08a | 0.83 ± 0.03b | 1.04 ± 0.05ab | 1.23 ± 0.09a | 1.29 ± 0.09a | 1.28 ± 0.06a |
| Internode 5 | 1.38 ± 0.07a | 0.85 ± 0.04b | 1.27 ± 0.05a | 1.44 ± 0.04a | 1.49 ± 0.06a | 1.52 ± 0.07a |
| **Internode diameter (cm)** | | | | | | |
| Internode 4 | 0.60 ± 0.02c | 0.61 ± 0.01c | 0.84 ± 0.02a | 0.74 ± 0.02b | 0.71 ± 0.01b | 0.68 ± 0.01b |
| Internode 5 | 0.59 ± 0.02c | 0.65 ± 0.02bc | 0.79 ± 0.02a | 0.70 ± 0.02b | 0.66 ± 0.01bc | 0.64 ± 0.02bc |
| Stem diameter (cm) | 0.60 ± 0.01ab | 0.61 ± 0.01ab | 0.65 ± 0.02a | 0.58 ± 0.01b | 0.57 ± 0.01b | 0.56 ± 0.01b |
| **Leaf insertion angle (°)** | | | | | | |
| Leaf 4 | 69 ± 1.18a | 62 ± 1.88a | 62 ± 1.25a | 62 ± 3.03a | 63 ± 1.90a | 70 ± 2.12a |
| Leaf 5 | 68 ± 2.14ab | 60 ± 1.32bc | 56 ± 1.67c | 61 ± 2.09bc | 67 ± 2.03ab | 70 ± 3.05a |
| **Reproductive characterization** | | | | | | |
| Sepals number | 5 ± 0.04a | 5 ± 0.02a | 5 ± 0.09a | 5 ± 0.08a | 5 ± 0.06a | 5 ± 0.07a |
| Petals number | 5 ± 0.03a | 5 ± 0.03a | 5 ± 0.06a | 5 ± 0.10a | 5 ± 0.08a | 5 ± 0.07a |
| Flowers of $1^{st}$ inf. | 7.7 ± 0.27c | 12.6 ± 0.28a | 9.5 ± 0.31b | 7.9 ± 0.21c | 8.6 ± 0.20bc | 8.4 ± 0.20c |
| Length of $1^{st}$ inf. | 10.04 ± 0.30b | 12.00 ± 0.38a | 11.95 ± 0.26a | 9.48 ± 0.24b | 9.60 ± 0.18b | 9.49 ± 0.13b |

Measurements were performed 60-days-after-germination for vegetative characterization and 20-days-after-flowering for reproductive characterization. Data are means ± SE n = 10. Different letters indicate statistically significant differences (Tukey's test, $P < 0.05$). inf. = inflorescence.

### Supplementary Fig. S1

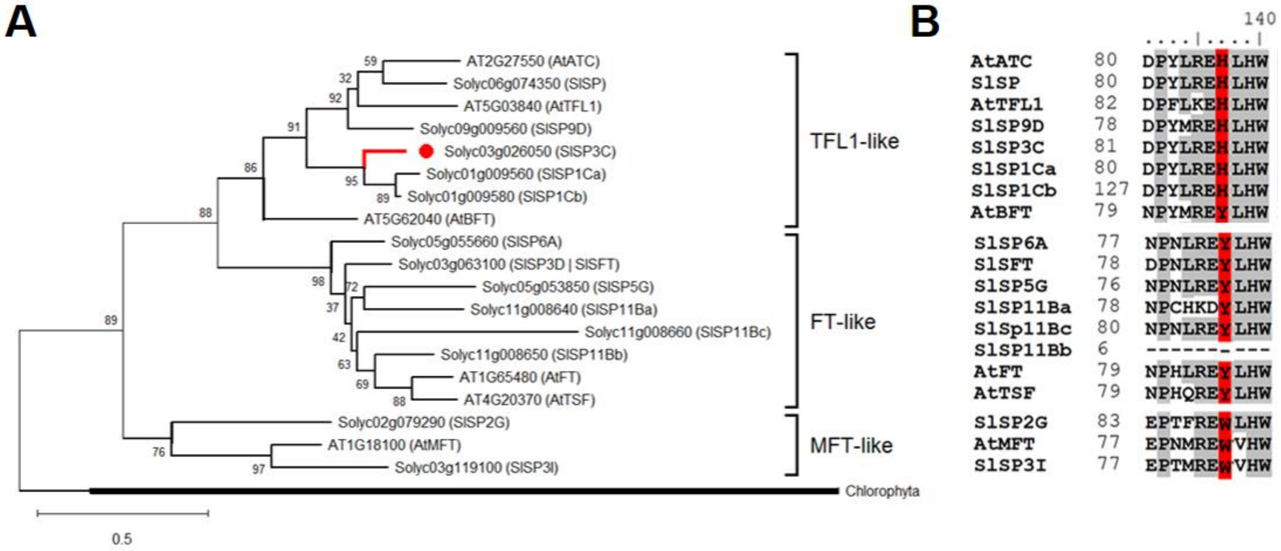


**Figure S1. Characterization of the *SP3C* gene.** (A) Phylogenetic reconstruction obtained from the alignment of *Arabidopsis thaliana*, tomato (*Solanum lycopersicum*) and Chlorophyta CETS (CENTRORADIALIS/TERMINAL FLOWER 1/SELF-PRUNING)-like proteins. All sequences are listed in Supplementary Table S1. (B) Detail of the alignment of *A. thaliana* and tomato CETS-like protein highlighting (red) the key amino acid for flowering repression (histidine-H) or activation (tyrosine-Y) activity (Hanzawa *et al.*, 2005). Positions with at least 50% residue identity are shaded in gray.

### Supplementary Fig. S2

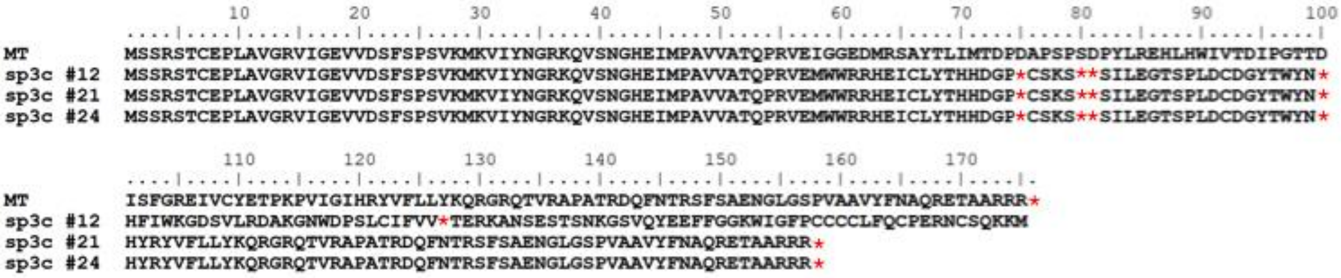


**Figure S2. Amino acid sequences of the *sp3c* mutant alleles.** The alignment of amino acid sequences compares the Micro-tom (MT) wild-type and three mutants generated by CRISPR/Cas9: *sp3c*#12, *sp3c*#21 and *sp3c*#24. The generated mutations led to the addition of premature stop codons in the protein of all three lines, resulting in predicted proteins of 74 amino acids. Stops codons positions are indicated by red asterisks.

**Supplementary Fig. S3**

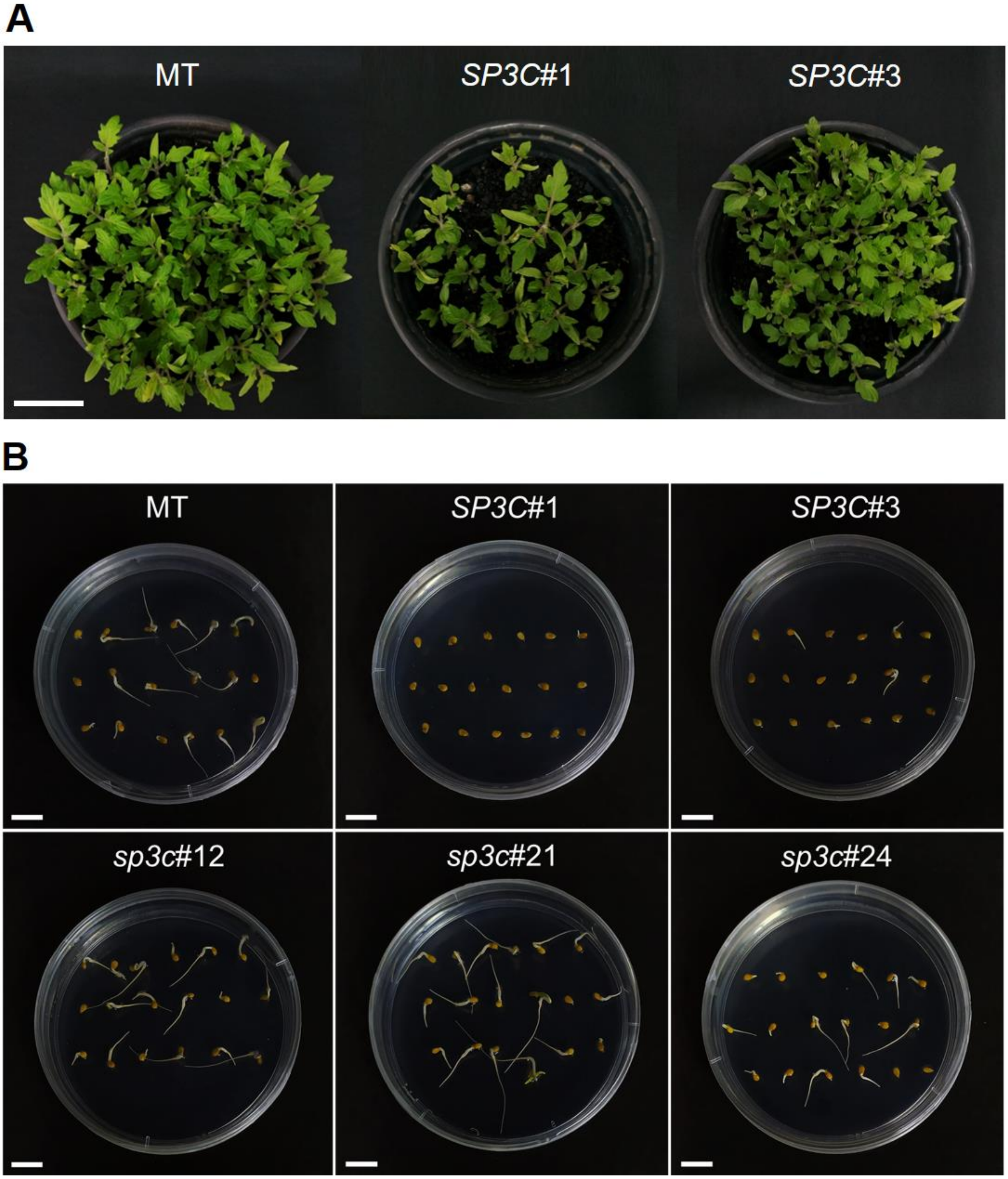


**Figure S3. *SP3C* acts as a repressor of seed germination.** (A) Differences between germination of Micro-tom (MT), *35S::SP3C*#1 (*SP3C*#1) and *35S::SP3C*#3 (*SP3C*#3) overexpression lines at 40-days-old. A total of 50 seeds were sown in each pot. Scale bars, 5 cm. (B) Representative images of germination assay made in MT, overexpression lines (*SP3C*#1; *SP3C*#3) and mutants (*sp3c*#12; *sp3c*#21; *sp3c*#24) at 5 days after sowing. The number of seeds with emitted root was evaluated daily. n = 4 plates per genotype with 18 seeds each. Scale bars, 1 cm.

**<u>Supplementary Fig. S4</u>**

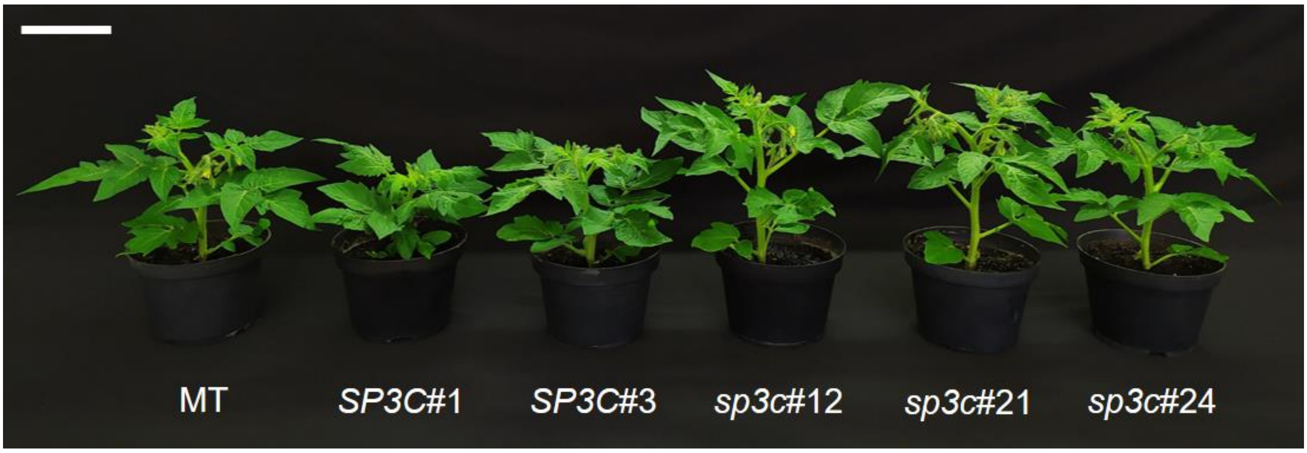


**Figure S4. Tomato plants harboring allelic variations in the *SP3C* gene.** Representative plants of *Solanum lycopersicum* cv. Micro-tom (MT), overexpression lines (*SP3C*#1 and *SP3C*#3) and mutants (*sp3c*#12, *sp3c*#21 and *sp3c*#24) with 36 days after sowing. Scale bars, 4 cm.

Supplementary Fig. S5

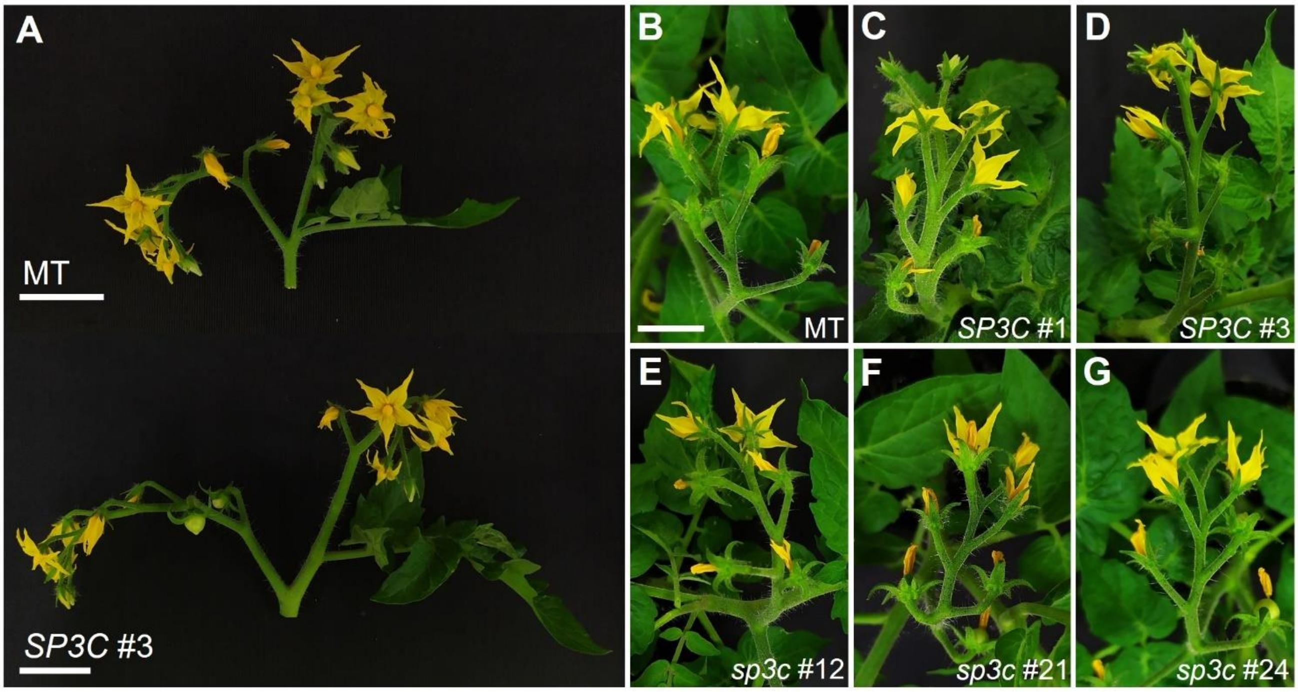


**Figure S5. High levels of *SP3C* increase the flowers number and the inflorescence length.** (A) Primary and secondary inflorescence of Micro-tom (MT) wild-type and overexpression line *SP3C*#3, at 20-days-after-flowering. (B-D) Inflorescence phenotype in MT, *35S::SP3C*#1 (*SP3C*#1) and *35S::SP3C*#3 (*SP3C*#3) overexpression lines, showing a longer inflorescence carrying a greater number of flowers. (E-G) Wild-type inflorescence phenotype in all *sp3c* mutant alleles. Scale bars, 2 cm.

**Supplementary Fig. S6**

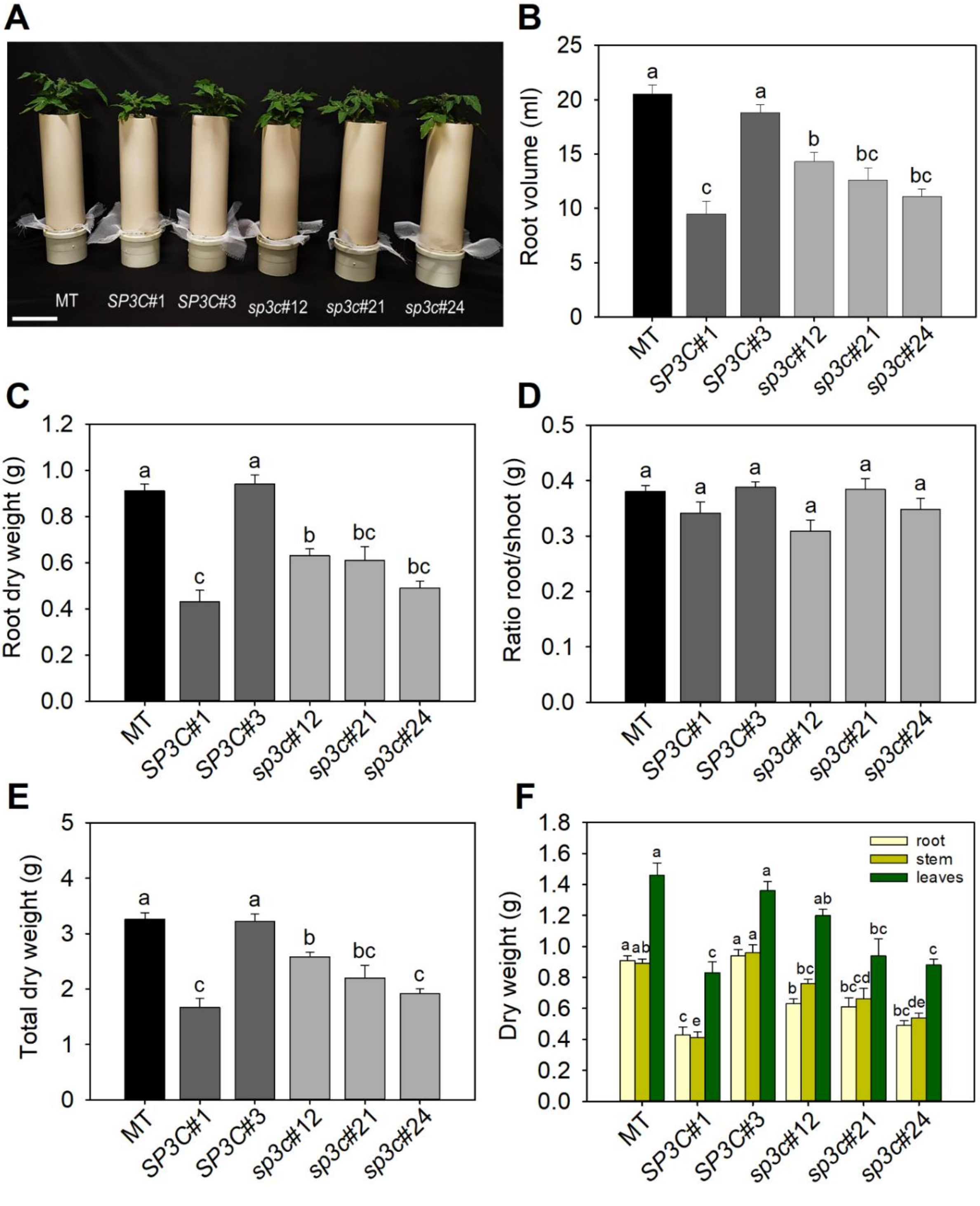


**Figure S6. Impacts of *SP3C* on root grown and development.** (A) Phenotype representative of Micro-Tom (MT), overexpression lines (*SP3C*#1 and *SP3C*#3) and mutants (*sp3c*#12, *sp3c*#21 and *sp3c*#24) at 50-days-old. The seedlings were grown in vertical PVC cylinders to track and quantify the root system development. Scale bars, 10 cm. (B) root volume, (C) root dry weight, (D) ratio root/shoot (with dry weight), (E) total dry weight, (F) dry weight of root, stem and leaves. Data are means ± SE (n = 10 plants). Different letters indicate statistically significant differences (ANOVA + Tukey's test, P <0.05).

**Supplementary Fig. S7**

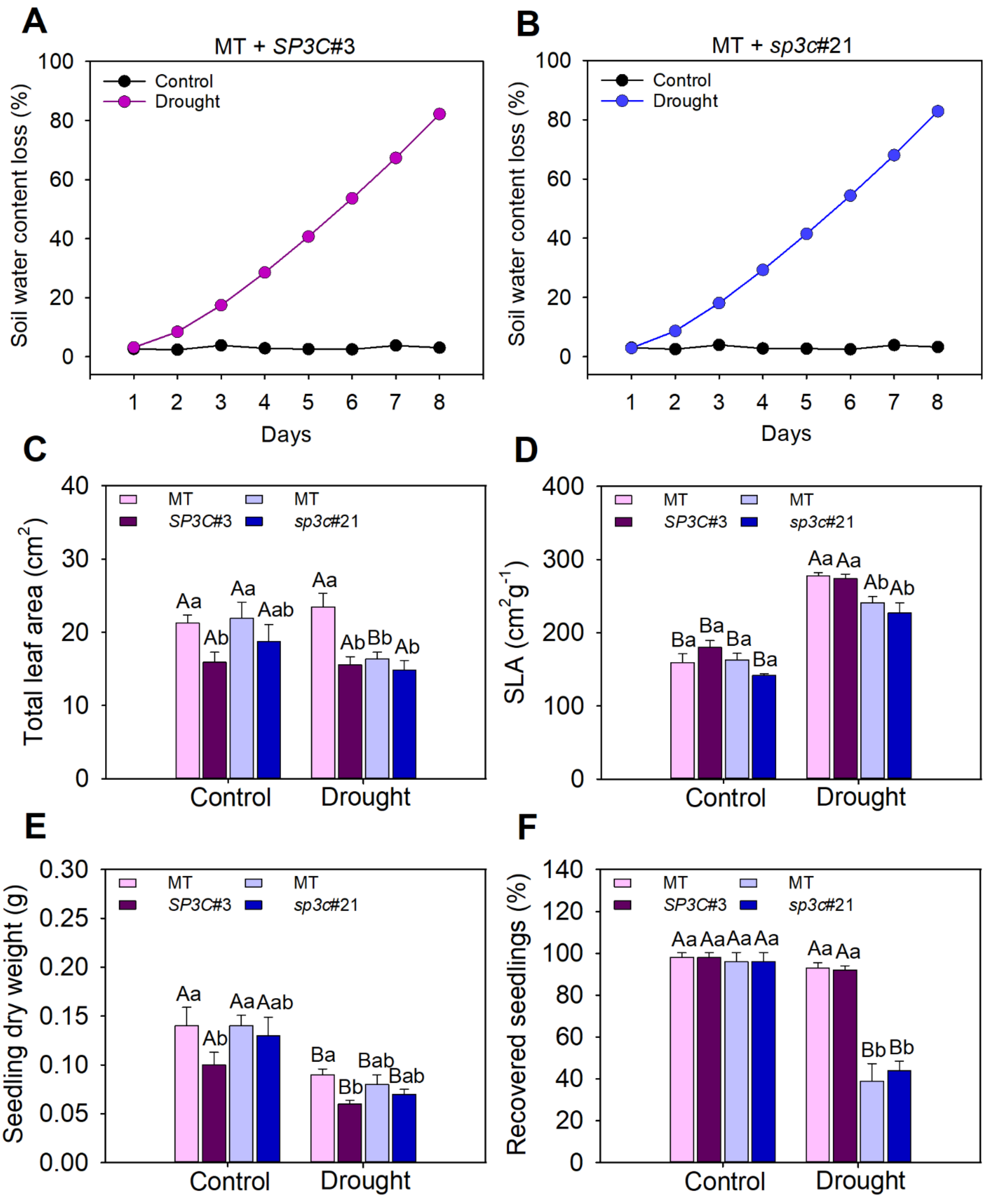


**Figure S7. *SP3C* improves tomato seedling response to drought.** (A and B) Water loss of soil measured during the 8 days of experiment, obtained by weighing of the pots. The pots under control condition were irrigated every day until the soil reaches its field capacity. (C) Total leaf area ($cm^2$) measured in MT/*SP3C*-overexpression (in purple) and MT/*sp3c*-mutant (in blue) respectively; (D) specific leaf area (SLA); (E) seedling dry weight (g) and (F) percentage of seedlings recovered after stress. Assessments were performed at the end of water stress, in 37-days-old seedlings. Data are means ± SE ($n \geq 4$). Different letters indicate statistically significant differences (Tukey's test, $P < 0.05$).
